# Noncoaxial Transcatheter Aortic Valve Deployment Creates Cusp-Specific Thrombogenic Microenvironments Through Altered Sinus Hemodynamics

**DOI:** 10.64898/2026.04.17.719323

**Authors:** Thangam Natarajan, Jae Hyun Kim, Christian Salgado, Akshita Jha, Cole Baker, Stephanie L. Sellers, Joseph Aslan, Monica Hinds, Ajit P. Yoganathan, Lakshmi Prasad Dasi

## Abstract

**Background:** Transcatheter aortic valve replacement has transformed the management of aortic stenosis; however, adverse outcomes such as leaflet thrombosis and hypoattenuating leaflet thickening remain clinically significant concerns. Flow disturbances resulting from valve canting may alter local hemodynamics and promote thrombogenic conditions. We investigated how modest transcatheter heart valve canting alters cusp-specific sinus flow and washout and promotes localized thrombogenic microenvironments associated with leaflet surface thrombus formation using particle image velocimetry, a physiologic blood loop, and tissue analysis.

**Methods:** A patient-derived aortic root model was used to evaluate the hemodynamic and thrombogenic effects of THV canting at -10° (anti-curvature), 0° (neutral), and +10° (along-curvature). High-resolution particle image velocimetry quantified sinus flow fields and washout characteristics, and complementary whole-blood loop experiments enabled histologic assessment of leaflet-associated thrombus formation.

**Results:** Canting redistributed systolic jet orientation and sinus recirculation in a direction-dependent manner while preserving global hemodynamic measurements. The most spatially constrained cusp showed the largest increase in stasis and the slowest washout. In the right coronary cusp, anti-curvature canting increased the fraction of sinus area with velocity magnitude <0.05 m/s to 92% versus 43% in neutral and 10% in along-curvature deployments, and prolonged neo-sinus (T_90_) washout to 4.7 cycles versus 2.9 and 1.8 cycles, respectively. Histology localized surface-adherent platelet/fibrin thrombus to these poorly washed regions, most prominently on the right coronary cusp leaflet in anti-curvature deployments. Left and noncoronary cusp responses shifted with tilt direction, indicating redistribution rather than uniform worsening of thrombogenic conditions.

**Conclusions:** Even modest noncoaxial deployment is sufficient to create sinus-resolved thrombogenic microenvironments that are not captured by global gradient or effective orifice area. Deployment configuration is therefore a modifiable determinant of post-TAVR leaflet thrombosis risk and may contribute to HALT.

## 1 Introduction

Transcatheter aortic valve replacement (TAVR) has become the standard therapy for severe aortic stenosis across a broad spectrum of patients, including younger and lower-risk populations. Although clinical outcomes are comparable to surgical bioprosthetic valves, transcatheter heart valves (THVs) remain susceptible to leaflet thrombosis, typically characterized by thrombus formation on the surface of one or more prosthetic leaflets. This phenomenon which may be subclinical or clinically overt, has emerged as an important concern for THV durability and safety.^1^ The reported incidence of subclinical leaflet thrombosis, with or without reduced leaflet motion, ranges from 7% to 35% across valve types.^2–4^ Potential consequences include thromboembolic events, stroke, and possibly premature structural valve degeneration.^5^ As TAVR expands to younger and lower-risk populations, understanding mechanisms that lead to leaflet thrombosis, and by extension, affect the structural integrity of the valve, has become increasingly important. Like obstructive valve thrombosis, THV structural deterioration arises from prothrombotic blood-cell responses that promote calcification, fibrin deposition, and leaflet degeneration, processes that can be mitigated by anticoagulation.^6–8^

A central mechanistic hypothesis is that abnormal sinus flow and localized blood stasis contribute to THV leaflet thrombosis.^9–11^ The aortic sinuses and neo-sinuses (the space between native leaflets and the transcatheter prosthesis) act as *“flow reservoirs”* where slow washout and low shear can favor clot formation. Computational simulations and benchtop models have supported this hypothesis and have shown that regions of low velocity, prolonged residence time, and reduced sinus washout are associated with increased thrombotic risk on bioprosthetic valves.^9,12,13^ These studies provide a plausible mechanistic link, i.e., local flow stasis and impaired transport; however, they do not clearly identify which procedural or geometric factors create adverse sinus environments in the first place.

Recent clinical imaging studies suggest that structural changes or deformations during THV deployment may influence these sinus flow environments and thereby affect thrombosis risk. In clinical practice, THV deployment is not always perfectly coaxial: minor deviations in alignment arise from asymmetric annular calcification, uneven expansion, or procedural variability, leading to valve canting relative to the aortic root axis. Retrospective CT analyses, including reports by Fukui et al.^14^ and subsequent meta-analyses, have linked canted or deformed THVs and other deployment parameters with higher rates of leaflet thrombosis and HALT.^15–18^ These observations imply that structural changes of the THV during deployment may be an upstream determinant of leaflet thrombosis. Yet, the mechanistic basis for this association remains unknown. Most prior work has emphasized global metrics such as transvalvular gradients, effective orifice area, or leaflet motion, without resolving sinus-specific flow fields and directly correlating hemodynamic disturbances with histologic evidence of leaflet surface thrombus.^12,19^ Consequently, whether modest malalignment creates sinus-dependent prothrombotic microenvironments has not been mechanistically evaluated in a controlled experimental setting.

To address this gap, we evaluated how controlled THV canting (balloon-expandable) alters the overall hemodynamics and thrombus formation using a population-representative aortic root model. High-resolution particle image velocimetry (PIV) was used to quantify sinus-specific flow patterns and transport characteristics. Under identical geometric and flow conditions, the working fluid was then replaced with porcine whole blood in a physiologic closed-loop system to enable leaflet histologic analysis and localization of surface-adherent thrombus.

In this study, we hypothesized that modest THV canting would generate cusp-specific sinus flow disturbances, prolong washout, and create localized prothrombotic microenvironments that promote thrombus formation on the leaflet surface. By integrating high-resolution flow mapping with transport analysis and histology, this study directly links THV canting to sinus hemodynamics and leaflet surface–level thrombi.

## 2 Methods

### 2.1 Patient-averaged Aortic root model

A patient-averaged aortic root geometry was generated from computed tomography angiography (CTA) datasets of 45 patients who underwent implantation of a 23-mm balloon-expandable transcatheter aortic valve for severe aortic stenosis. Patient selection criteria and imaging acquisition details have been reported previously.^20^ CTA images were segmented to reconstruct three-dimensional (3D) surface geometries of the aortic root. The resulting surface meshes were smoothed and imported into a statistical shape modeling framework (ShapeWorks), which employs a particle-based entropy-minimization algorithm to establish point correspondences across anatomical surfaces and to derive the population mean geometry.^21^ The resulting mean aortic root shape was further smoothed to remove minor surface artifacts and used as the reference geometry for chamber fabrication. To fabricate the experimental chamber, the mean geometry was 3D-printed using stereolithography (Formlabs Form 3B, White Resin) to produce molds. Optically transparent polydimethylsiloxane (PDMS) chambers were then produced using a 10:1 base-to-curing-agent mixture (Sylgard™ 184 Silicone Elastomer Kit, Dow, CA, USA). The mixture was degassed under vacuum to remove entrapped air bubbles and poured around the printed mold. PDMS was cured at 60 °C for 4 hours, followed by 24 hours at room temperature to complete polymerization. The optically transparent PDMS chamber enabled both high-resolution PIV measurements and subsequent whole-blood circulation experiments.

### 2.2 Custom THV Design and Fabrication

Custom THVs, structurally and functionally similar to commercial balloon expandable valves, were constructed using glutaraldehyde-fixed porcine pericardial tissue sheets (PeriSeal™ Tissue Patch, Avalon Medical, Stillwater, MN, USA). Leaflet patterns were laser-cut from 10 × 10 cm pericardial sheets, using templates designed to achieve a 23-mm valve diameter. The cut leaflets were sutured using 7-0 polypropylene suture to form individual cusps in a two-dimensional configuration. These cusps were subsequently joined along the commissural edges to create the three-dimensional leaflet structure. The assembled leaflets were then sutured at the base and commissures to a 23-mm laser-cut cobalt-chromium stent frame geometrically modeled after the SAPIEN 3 transcatheter valve (supplemental figure S1 A–C). To characterize the surface morphology of the pericardial tissue, representative samples (1 × 1 cm) were prepared for scanning electron microscopy (SEM). Samples were dehydrated through a graded isopropyl alcohol series, air dried, and sputter-coated with gold (Cressington 108) prior to imaging with a Phenom XL G2 scanning electron microscope at an accelerating voltage of 15 kV (supplemental figure S1 D–E).

### 2.3 THV Deployment and Canting Protocol

Three deployment configurations were evaluated (Figure 1): (1) a -10^°^ tilt toward the native right-coronary cusp (RCC) referred to as anti-curvature, (2) a neutral control (0^°^ tilt), and (3) a +10^°^ tilt away from the RCC, referred to as along-curvature. Tilting or canting was defined as the angular deviation of the valve longitudinal axis relative to the normal of the annular plane. Under anti-curvature canting (-10^°^), the RCC functioned as the primary space-constrained cusp, whereas the non-coronary cusp (NCC) and left coronary cusp (LCC) retained relatively larger sinus regions. In contrast, under along-curvature canting (+10^°^), the NCC and LCC experienced partial spatial constraint while the RCC remained relatively unconstrained. To impose controlled canting in the experiments, a custom 3D-printed tapered annular disk was fabricated with a fixed 10^°^ inclination relative to its base plane (see supplemental figure S2). The transcatheter valve was sutured circumferentially onto the tapered surface, ensuring its longitudinal axis aligned with the imposed incline. The valve-disk assembly was then deployed into the aortic root model while maintaining a constant implantation depth and radial expansion across all tested configurations. This approach ensured that the prescribed canting angle was geometrically imposed and preserved during experimentation and served as the primary independent variable influencing sinus hemodynamics.

**Figure 1.**
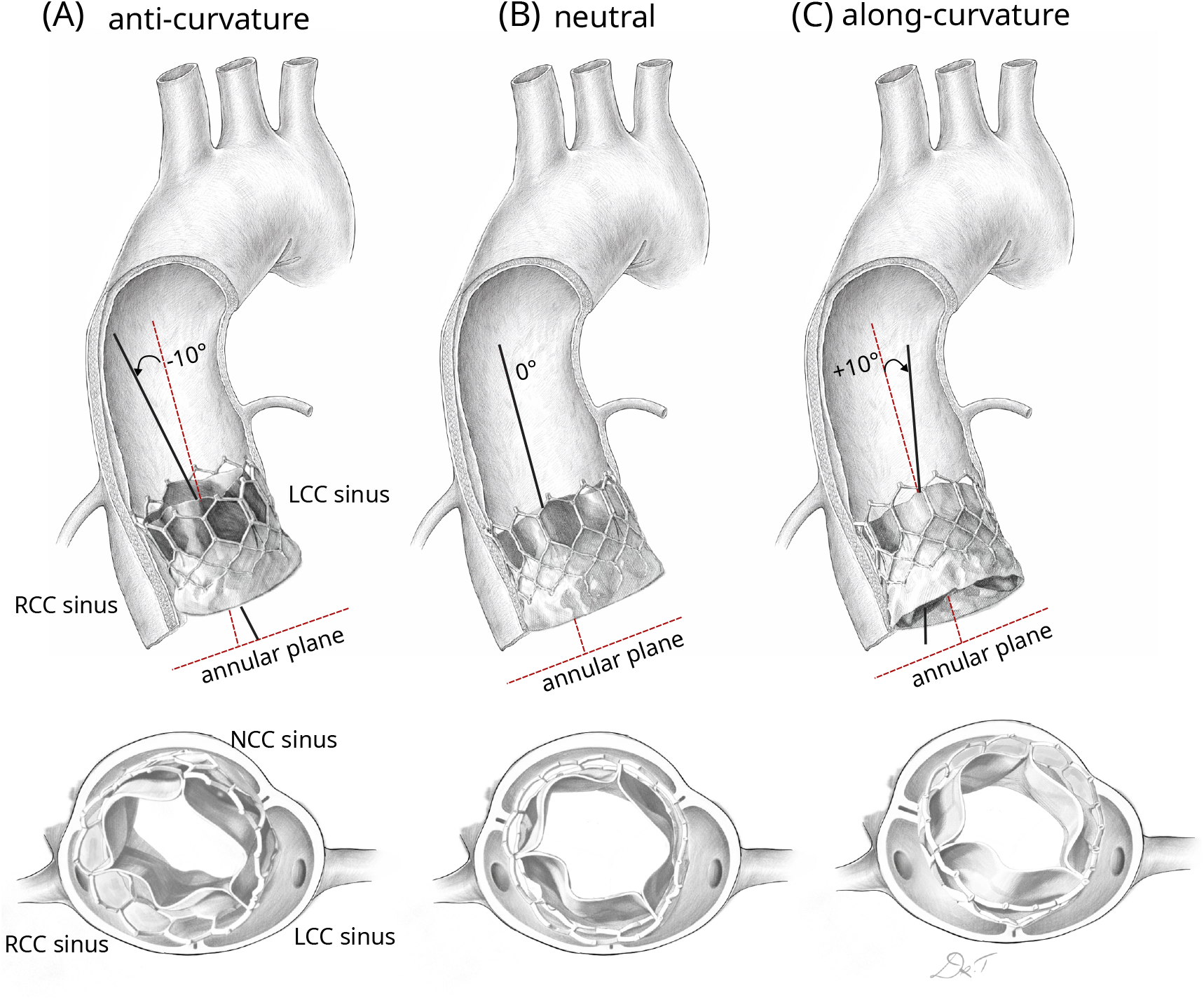
Schematic of THV canting configurations in the patient-derived aortic root model. (A) -10^°^ canting of the THV axis toward the right coronary sinus relative to the annular plane normal. (B) Neutral deployment (0^°^), with the THV axis aligned with the root axis. (C) +10^°^ canting of the THV axis toward the left and non-coronary sinus relative to the annular plane normal. The bottom panel shows the top view of the sinus of Valsalva with the different THV configurations canted with respect to the sinuses.

### 2.4 In Vitro Circulatory Loop for PIV

Leaflet kinematics and flow fields were evaluated using the Georgia Tech Left Heart Simulator, a pulsatile flow loop that reproduces physiologic cardiac conditions.^15,17^ The schematic of the experimental setup is shown in supplemental figure S3 A. Experiments were conducted under nominal physiologic parameters (cardiac output 3.5 L/min, stroke volume 60 mL, mean arterial pressure 100 mmHg). A water-glycerin blood analog (36% glycerin by volume; dynamic viscosity ≈ 3.5 cP) was used to match the density and viscosity of blood while maintaining optical transparency for flow measurements. Planar particle image velocimetry (PIV) was used to quantify velocity fields within the sinus, neo-sinus, and downstream regions of the THV. The flow was seeded with fluorescent tracer particles (1-20 *µ*m diameter). A pulsed laser generated an approximately 1-mm-thick light sheet aligned with the valve centerline, capturing one commissural post and the coaptation line between the leaflets within the imaging plane. The model chamber was rotated to interrogate each cusp region individually. Image pairs were acquired at 500 Hz using a high-speed camera system synchronized to a trigger. For each canting configuration, image acquisition was performed during defined phases of systole (acceleration, peak systole, and deceleration), and ensemble averaging over 20 cardiac cycles was used to obtain phase-resolved velocity fields. Velocity vectors were calculated using cross-correlation algorithms in DaVis (LaVision). Vorticity magnitudes (*ω*) were derived from spatial gradients of the in-plane velocity components to quantify local rotational flow structures. Sinus washout was evaluated using Lagrangian particle tracking applied to the phase-resolved velocity fields. Virtual particles were uniformly seeded within predefined sinus and neo-sinus regions of interest and advected using temporally and spatially interpolated velocity data. Washout time was defined as the time required for 90% (T_90_) of particles to exit the region, enabling normalized comparison of sinus transport dynamics across deployment configurations.

### 2.5 In Vitro Circulatory Loop for Whole Blood

Whole-blood experiments were conducted using a modified pulsatile circulatory loop designed to accommodate hemocompatible operation while preserving physiologic flow conditions.^22^ The schematic of the whole blood circulatory loop is shown in supplemental figure S3 B. The system consisted of a closed-loop pulsatile flow circuit driven by a computer-controlled linear actuator coupled to a piston pump, generating physiologic aortic pressure and flow waveforms across the test section. The PDMS aortic root chamber containing the THV was mounted within the loop, with the circulating fluid replaced by porcine whole blood. System compliance and resistance elements were adjusted to reproduce target physiologic parameters, including cardiac output, stroke volume, heart rate, and systolic and diastolic pressures. Flow rate was measured with an inline flow probe downstream of the test section, while high-fidelity pressure transducers recorded ventricular- and aortic-side pressures throughout the cardiac cycle. Signals were acquired using a synchronized data-acquisition system, and measurements were averaged over multiple cardiac cycles after stable periodic operation was established. Transvalvular pressure gradients and effective orifice areas were calculated from simultaneous flow and pressure measurements to confirm physiologic valve performance. All other parameters and conditions were kept identical to the Georgia Tech left heart simulator described above, and the only change was the circulating fluid, which is porcine whole blood.

### 2.6 Whole Blood Circulatory Loop Protocol

Fresh porcine whole blood anticoagulated with 3.2% sodium citrate was obtained from a commercial vendor (Lampire Biological Laboratories, Pipersville, PA, USA) following same-day collection and shipment. Experiments were conducted in citrated whole blood without recalcification in order to preferentially interrogate platelet-mediated responses to altered sinus hemodynamics while limiting systemic (loop-specific) coagulation cascade amplification. Prior to each experiment, whole blood was maintained under low-shear orbital agitation using an orbital shaker (Belly Dancer, Stovall Life Science, Inc., Greensboro, NC, USA) to prevent cellular sedimentation without inducing preactivation at room temperature (25-30 ^°^C). Baseline hematologic parameters, including platelet count and activation status, were measured by flow cytometry to confirm consistency across experimental runs. The pulsatile flow loop was maintained at room temperature under physiologic cardiac output and heart rate conditions. After each run, blood samples were collected for flow cytometric quantification of platelet activation, and valve leaflets were harvested for histologic assessment of surface thrombus deposition.

### 2.7 Histologic and Flow Cytometry Analysis

The THVs from the whole-blood experiments were fixed and stored in 10% (v/v) neutral buffered formalin for further histological and immunohistochemical analyses. The leaflets were then removed from the stent frame, bisected, and cut into 2–4 mm sections and embedded in paraffin, resulting in 5 tissue cross sections analyzed per leaflet. Paraffin-embedded leaflet sections (5 *µ*m) were deparaffinized, rehydrated, and stained using Carstairs’ method to distinguish platelets (gray-blue to navy) from fibrin (red), with collagen appearing bright blue. High-resolution images of Carstairs-stained leaflet sections were acquired using a Zeiss Axio Observer Z1 microscope. For flow cytometry analysis, whole blood was collected from the in vitro loop using 1 mL Luer-Lock tip syringes and diluted 1:4 with HEPES-buffered Tyrode’s solution. Diluted whole blood was mixed with bovine lactadherin-FITC (Prolytix, Essex Junction, VT), fluorescent fibrinogen Alexa-488 (ThermoFisher, Waltham, MA), CD61-Alexa647 (JM2E5, Bio-Rad, Hercules, CA), and/or CD62P-647 (Psel.KO.2.7, Bio-Rad, Hercules, CA) for 30 minutes at 37 ^°^C. Positive control tubes had added phorbol 12-myristate 13-acetate (PMA, 5 *µ*m) to induce platelet activation. After incubation, samples were fixed with 4% paraformaldehyde, wrapped in parafilm, and stored at 4 ^°^C. Samples were then analyzed on a FACSCanto II flow cytometer (Becton-Dickinson, Franklin Lakes, NJ, USA). Populations were gated according to forward scatter (FSC-A) and side scatter (SSC-A) to distinguish platelets, and CD61 fluorescence was used to verify platelet identity. Flow cytometry data were processed and analyzed using FlowJo software (version 10.7.1).

## 3 Results

### 3.1 Global Hemodynamics

Controlled canting of the THV altered the spatial relationship between the valve frame and the aortic sinuses, producing cusp-specific differences in sinus flow patterns and thrombus deposition (Figures 2–8 and supplemental video). Introducing valve canting produced eccentric systolic jets and asymmetric shear layers that varied with the direction of canting relative to the aortic curvature. These changes altered the local flow environment within individual sinus cavities, resulting in cusp-specific differences in velocity magnitude, vortex formation, and sinus recirculation patterns. The effective orifice area (EOA) and mean transvalvular pressure gradient (TVPG) (see supplemental figure S4 A-B) are comparable to those of a commercial SAPIEN valve, and the intra-configuration differences are negligible. The downstream and cusp-specific flow characteristics and associated leaflet histology are described below.

**Figure 2.**
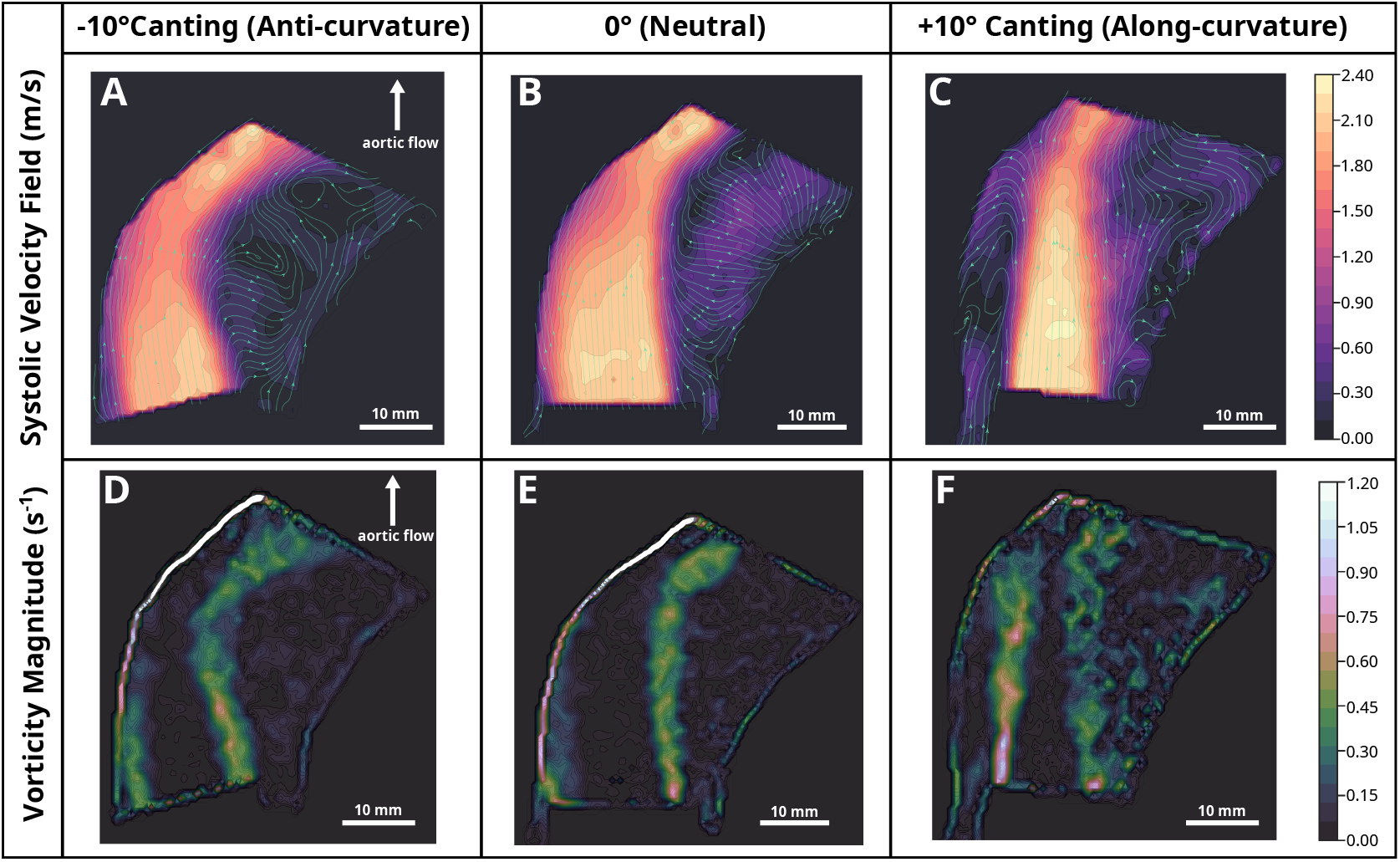
Effect of THV canting on downstream flow patterns. Panels A–C show systolic velocity vector fields within the downstream plane for anti-curvature (-10^°^), neutral (0^°^), and along-curvature (+10^°^) canting, respectively. Panels D–F display corresponding vorticity magnitude.

### 3.2 Downstream Jet and Vorticity Characteristics

Peak-systolic downstream velocity and vorticity fields associated with the three valve configurations are shown in Figure 2. Transvalvular pressure gradient and peak jet velocity were comparable across all configurations. In the neutral configuration (0^°^), the systolic jet remained largely aligned with the valve centerline and impinged on the outer wall of the proximal ascending aorta. A shear-induced region of retrograde flow was confined primarily near the inner wall of the aorta. Anti-curvature canting (-10^°^) produced a more pronounced lateral displacement of the jet toward the outer wall. This configuration generated an expanded region of retrograde flow along the inner wall and a larger vortical structure extending toward the outer curvature. In contrast, along-curvature canting (+10^°^) resulted in a more centrally directed jet that produced two smaller retrograde flow regions adjacent to both the inner and outer aortic walls. Quantitative vorticity measurements demonstrated configuration-dependent differences in rotational flow structures. Vorticity magnitude near the outer wall of the ascending aorta was greatest in the along-curvature configuration (≈ 1000 s^−1^), followed by the neutral configuration (≈ 400 s^−1^), and was lowest in the anti-curvature configuration (300 s^−1^). In contrast, the vorticity near the inner wall was highest in the anti-curvature configuration (≈ 2000 s^−1^), followed by neutral configuration (≈ 1000 s^−1^), and lowest in the along-curvature configuration(≈ 700 s^−1^). Higher vorticity magnitudes are commonly associated with larger recirculation zones, which promote retrograde flow into the corresponding aortic cusps.^23^ In the patient-averaged aortic root model in this study, the outer wall corresponds to the RCC, whereas the inner wall corresponds to the LCC. Reduced vorticity magnitude along the outer wall may limit the retrograde flow into the RCC, thereby increasing the likelihood of flow stasis, thrombosis, and fibrin deposition in this region. Similarly, lower vorticity near the inner wall may limit retrograde flow back into the LCC, thereby increasing the likelihood of flow stasis, thrombosis, and fibrin deposition on the corresponding leaflets.

### 3.3 Sinus Flow Stasis Distribution

The distribution of low-velocity regions within the aortic sinuses is summarized in Figure 3. Low-velocity regions were defined as areas with velocity magnitude |v| < 0.05 m/s, quantified from phase-averaged PIV velocity fields over the cardiac cycle. Under neutral canting (0^°^), low-velocity regions were relatively evenly distributed among the three sinuses. Anti-curvature canting (-10^°^) markedly increased the low-velocity area within the right coronary cusp (RCC) sinus, indicating greater flow stagnation in this region. In contrast, along-curvature canting (+10^°^) reduced the low-velocity area within the RCC while shifting regions of flow stasis toward the left and non-coronary cusps (LCC and NCC). For the RCC, the area exhibiting velocity magnitude below 0.05 m/s was lowest in the along-curvature configuration (10%), intermediate in the neutral configuration (43%), and highest in the anti-curvature configuration (92%). In the LCC, the percentage of sinus area with velocity magnitude below 0.05 m/s showed limited variation across configurations (40% for neutral, 50% for along-curvature, and 38% for anti-curvature). Finally, in the NCC, the percentage of the sinus area exhibiting velocity magnitude below 0.05 m/s was lowest in the anti-curvature configuration (11%), followed by the neutral configuration (42%) and the along-curvature configuration (52%). These results quantitatively demonstrate that valve canting redistributes sinus flow stagnation among individual cusps.

**Figure 3.**
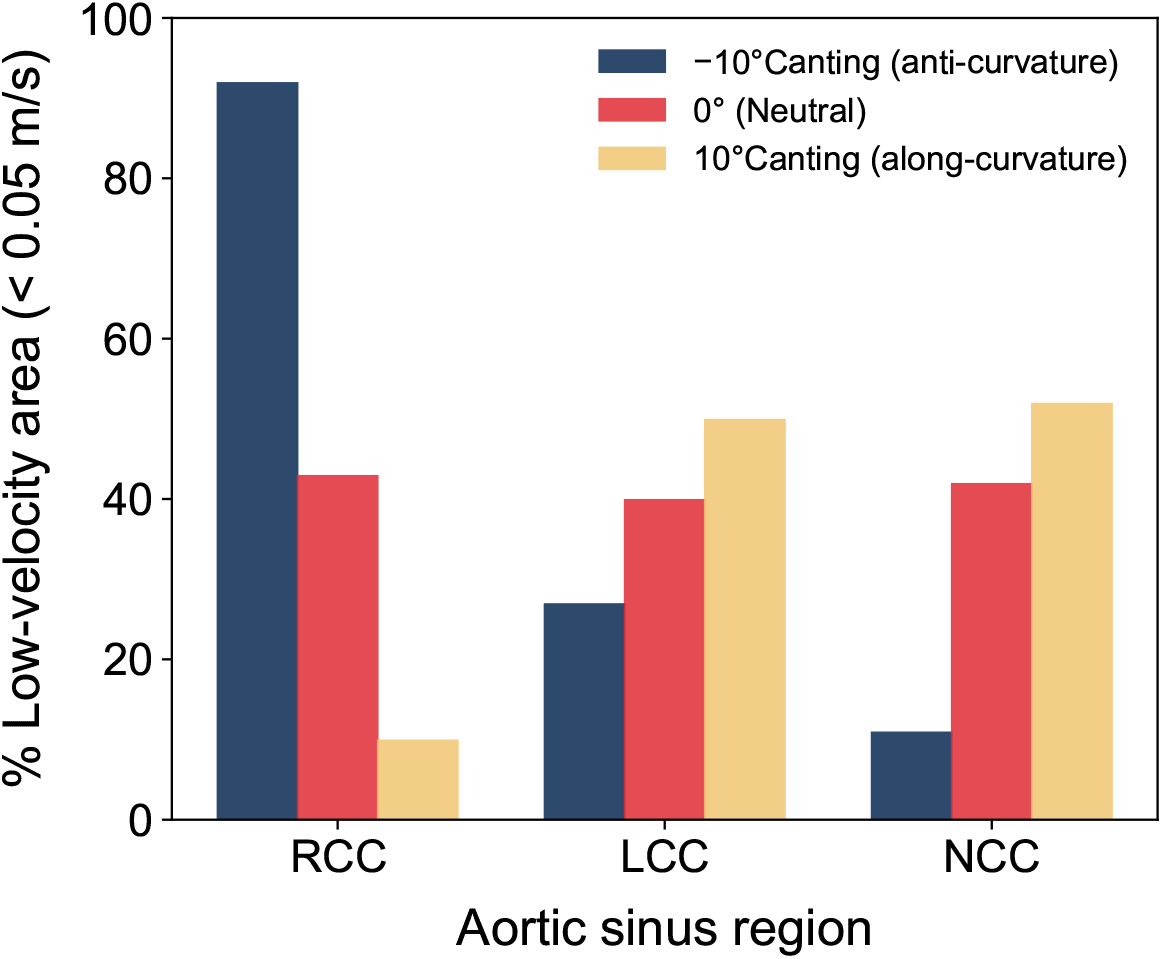
Percentage of aortic sinus area exhibiting low velocity (|v| < 0.05 m/s) quantified from phase-averaged PIV velocity fields averaged over x cardiac cycles. Anti-curvature canting (-10^°^) markedly increased low-velocity area within the right coronary cusp (RCC), whereas along-curvature canting (+10^°^) reduced RCC stasis and shifted low-velocity regions toward the left and non-coronary cusps (LCC and RCC). Neutral alignment (0^°^) produced a more uniform distribution across the three sinuses.

### 3.4 Platelet Integrity and Baseline Activation State

Flow cytometry analysis demonstrated preservation of platelet integrity and baseline activation state over the 4-hour circulation (Figure 4). Platelet populations remained stable based on FSC/SSC gating, with consistent CD41 expression at the beginning and end of the experiment, indicating minimal spontaneous platelet activation within the loop.

**Figure 4.**
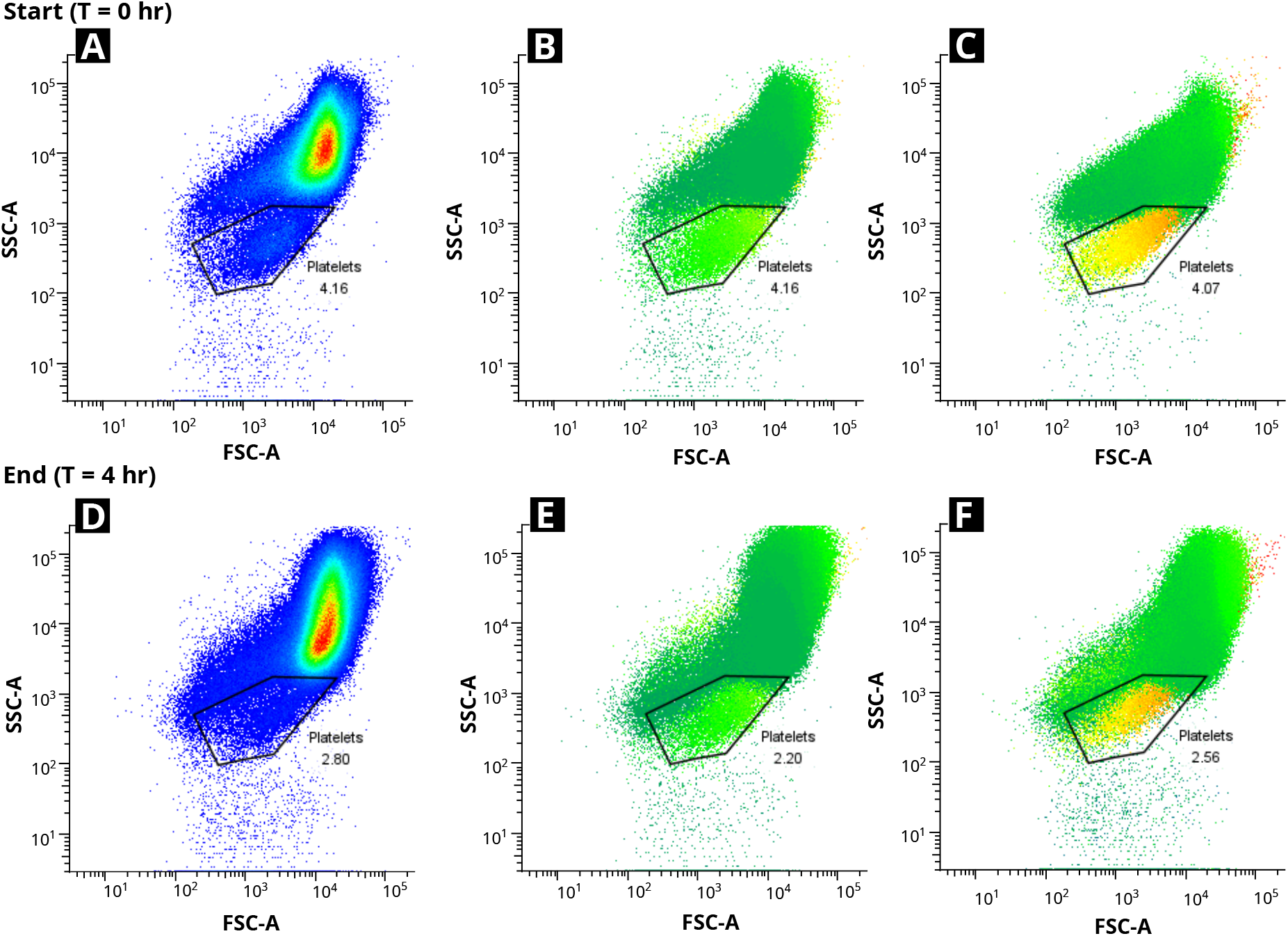
Flow cytometry analysis of porcine platelets in a 4-hour blood loop. FSC-A vs SSC-A plots (left) show the platelet population (gated), while CD41 overlay (right) confirms platelet identity. Top row: beginning of loop; bottom row: after 4 hours. Platelet percentages remained stable, indicating preserved platelet integrity.

### 3.5 Cusp-Specific Hemodynamics and Leaflet Thrombosis: Right Coronary Cusp

Representative velocity fields, vorticity distributions, and histological sections corresponding to the RCC are shown in Figure 5. During peak systole, leaflet opening was accompanied by increased velocity within the neo-sinus and localized recirculation into the native sinus. During flow deceleration and early diastole, leaflet closure produced retrograde flow into both the neo-sinus and native sinus. Among the three cusps, the RCC demonstrated the largest differences in flow characteristics across canting configurations. The along-curvature configuration exhibited the highest peak average velocity (0.15 m/s), followed by the neutral configuration (0.09 m/s) and the anti-curvature configuration (0.05 m/s). Histological sections of the RCC leaflet (Figure 5 G–I) showed extensive fibrin-rich surface thrombus on the aortic side of the leaflet in the anti-curvature configuration. Collagen fibers in this region appeared moderately disrupted compared with the other two configurations. In contrast, no visible fibrin or platelet aggregates were observed on the RCC leaflet in neutral or along-curvature configurations.

**Figure 5.**
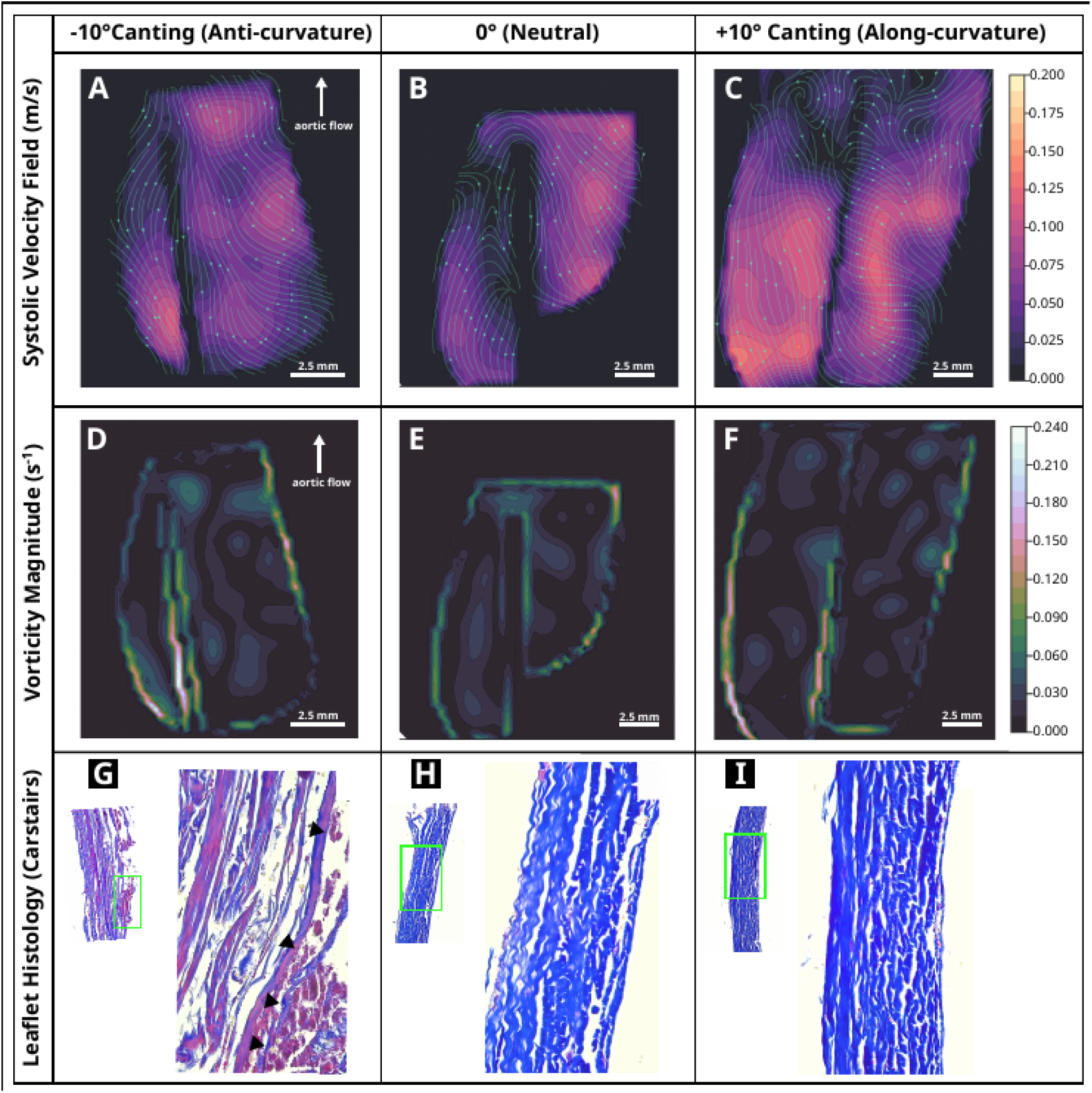
Effect of THV canting on RCC sinus flow patterns and RCC leaflet surface thrombosis. Panels A-C show systolic velocity vector fields within the RCC sinus plane for Anti-curvature (-10^°^), neutral (0^°^), and along-curvature (+10^°^) canting configurations, respectively. Panels D–F display corresponding vorticity magnitude maps highlighting enlarged shear layers and recirculation zones adjacent to the outflow leaflet belly under canted conditions. Panels G–I present representative Carstairs-stained histologic sections from the RCC leaflet outflow surface, with arrow heads indicating surface-adherent fibrin/platelet-rich thrombus that increases with canting, while underlying leaflet architecture remains preserved.

### 3.6 Cusp-Specific Hemodynamics and Leaflet Thrombosis: Left and Non-Coronary Cusp

The velocity fields and histological sections for the LCC and NCC are presented in Figure 6 and 7, respectively. Compared with the other cusps, the LCC exhibited smaller differences in flow patterns across the three canting configurations. Peak average velocity showed minimal variation between the neutral configuration (0.085 m/s), the along-curvature configuration (0.078 m/s), and the anti-curvature configuration (0.072 m/s). Corresponding histological sections (Figure 6 G–I) of the LCC leaflet demonstrated relatively limited thrombus deposition compared with the RCC. On observing the NCC sinus, the anti-curvature configuration exhibited the highest peak average sinus velocity (0.13 m/s), followed by the neutral configuration (0.09 m/s) and the along-curvature configuration (0.07 m/s). Histological sections of the NCC and LCC leaflets (Figure 6 and 7 G–I) showed intact, well-organized blue-staining collagen in all configurations, with no discrete throm-bus masses. In the anti-curvature configuration, a very faint, patchy pink hue was present along the aortic surface of the NCC leaflet, whereas no appreciable pink staining was observed in the neutral or along-curvature configurations.

**Figure 6.**
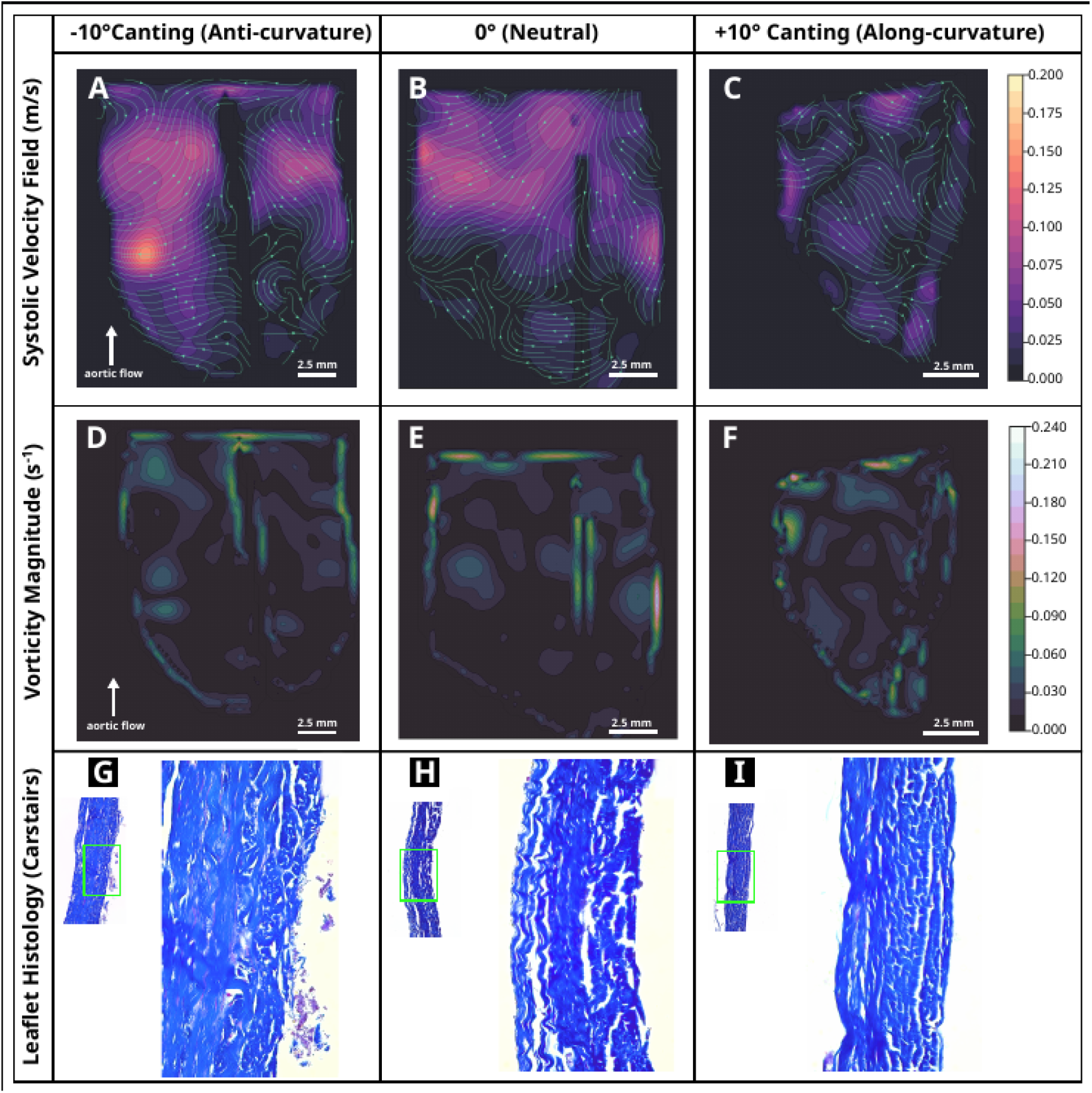
Effect of THV canting on LCC sinus flow patterns and LCC leaflet surface thrombosis. Panels A–C show systolic velocity vector fields within the LCC sinus plane for Anti-curvature (-10^°^), neutral (0^°^), and along-curvature (+10^°^) canting configurations, respectively. Panels D-F display corresponding vorticity magnitude maps highlighting enlarged shear layers and recirculation zones adjacent to the outflow leaflet belly under canted conditions. Panels G-I present representative Carstairs-stained histologic sections from the leaflet outflow surface, with arrow heads indicating surface-adherent fibrin/platelet-rich thrombus that increases with canting while underlying leaflet architecture remains preserved.

**Figure 7.**
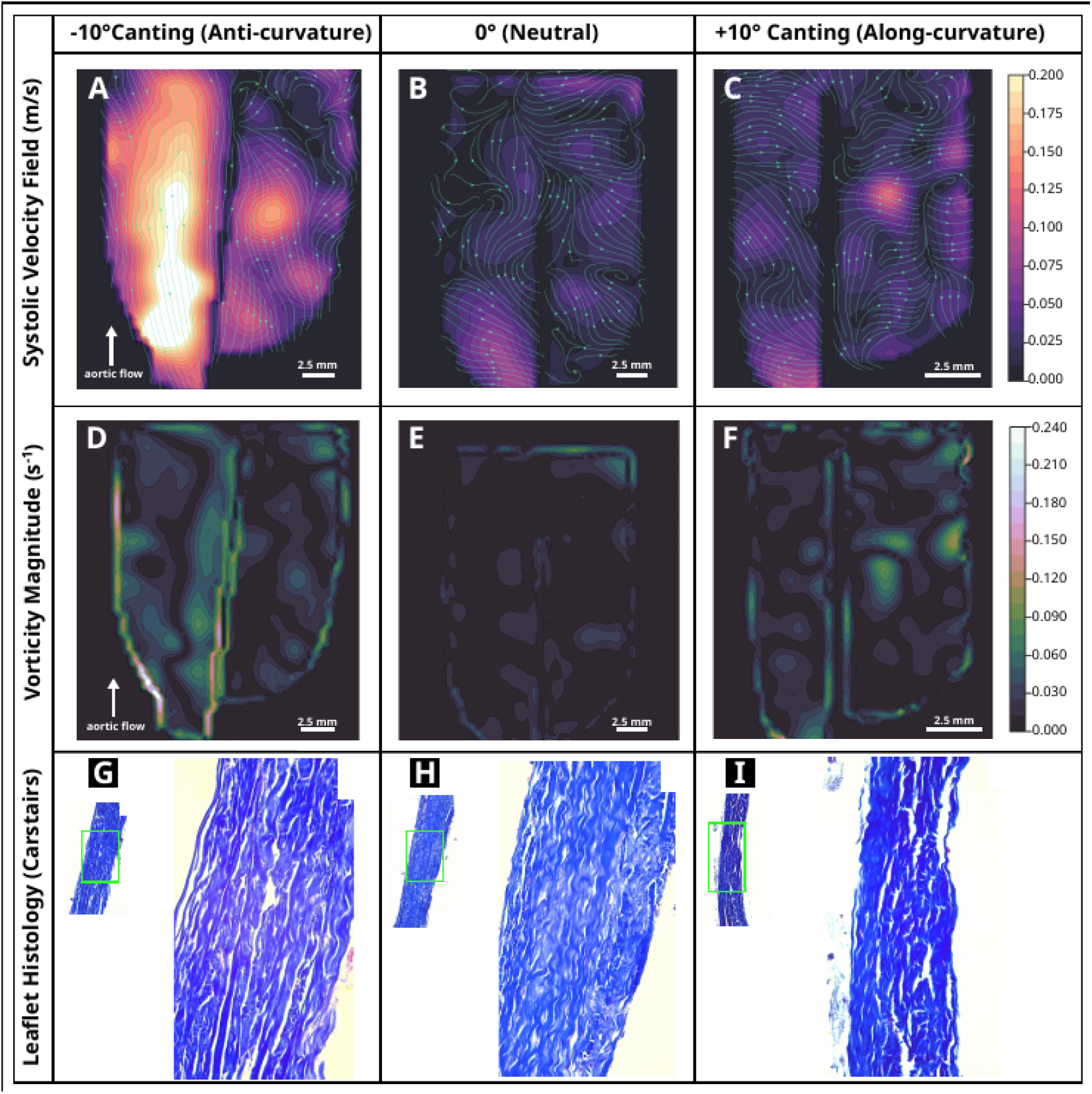
Effect of THV canting on NCC sinus flow patterns and NCC leaflet surface thrombosis. Panels A-C show systolic velocity vector fields within the NCC sinus plane for Anti-curvature (-10^°^), neutral (0^°^), and along-curvature (+10^°^) canting configurations, respectively. Panels D-F display corresponding vorticity magnitude maps highlighting enlarged shear layers and recirculation zones adjacent to the outflow leaflet belly under canted conditions. Panels G-I present representative Carstairs-stained histologic sections from the leaflet outflow surface, with arrow heads indicating surface-adherent fibrin/platelet-rich thrombus that increases with canting while underlying leaflet architecture remains preserved.

### 3.7 Neo-sinus Washout Characteristics

Figure 8 presents the particle residence curves as a percentage of particles remaining within the neo-sinus over time. More rapid decay of these curves to zero indicates faster washout within the neo-sinus region associated with the THV. Overall, the neo-sinus washout time showed a trend very similar to that of flow quantifications in the vicinity of the THV. In the RCC, washout time to reach 10% of particles remaining in the neo-sinus was the highest in the anti-curvature configuration (4.7 cycles), followed by the neutral configuration (2.9 cycles), and the along-curvature configuration (1.8 cycles). In contrast, in the NCC, the washout time to reach 10% of the particles remaining was longest in the along-curvature configuration (1.7 cycles), followed by the neutral configuration (1.5 cycles) and the anti-curvature configuration (0.6 cycles). Lastly, in the LCC, the washout time to reach 10% of the remaining particles was highest in the anti-curvature configuration (2.4 cycles), followed by the control configuration (1.6 cycles) and the along-tilt configuration (0.8 cycles).

**Figure 8.**
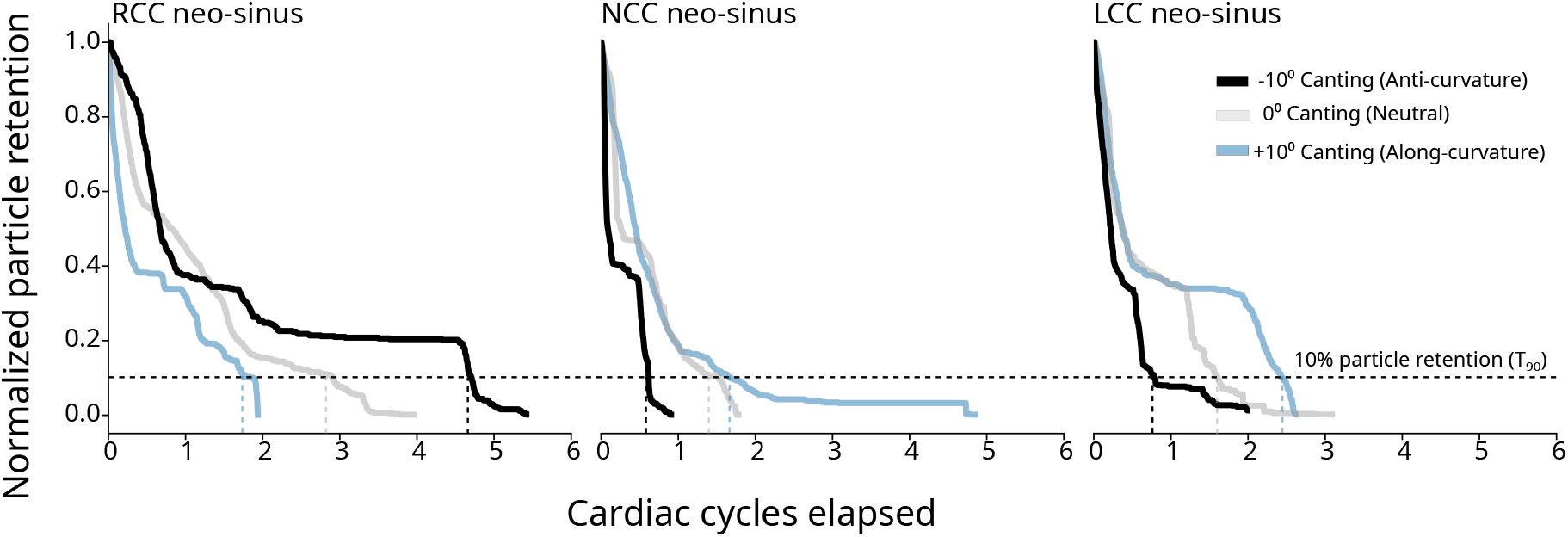
Sinus-specific washout dynamics across valve canting configurations. Normalized particle retention curves are shown for the right coronary cusp (RCC), non-coronary cusp (NCC), and left coronary cusp (LCC) neo-sinuses under anti-curvature (-10^°^), neutral (0^°^), and along-curvature (+10^°^) canting configurations. The horizontal dashed line denotes 10% particle retention, corresponding to the time to 90% washout (T_90_). Across all sinuses, anti-curvature canting consistently prolonged washout, evidenced by delayed T_90_ relative to neutral and along-curvature configurations.

## 4 Discussion

This study demonstrates that modest THV canting can redistribute sinus flow between individual aortic cusps, producing localized regions of stagnation that promote surface-level leaflet thrombosis despite similar EOA and TVPG across canting configurations. Using high-resolution PIV and a whole-blood circulatory loop, we show that canting angles of *±*10^°^ substantially reorganize downstream jet structure and sinus vortex patterns, leading to cusp-specific differences in flow stasis and washout characteristics. Regions exhibiting impaired sinus transport colocalized with Carstairs-positive thrombus on the fibrous leaflet surface. Importantly, these localized flow disturbances occurred without significant changes in global hemodynamic indices, such as effective orifice area or transvalvular gradient, indicating that conventional valve performance metrics may not capture the thrombogenic microenvironments created by noncoaxial deployment.

The aortic sinus functions as a flow-regulated reservoir in which physiologic vortices renew surface shear, facilitate particle clearance, and prevent prolonged stagnation near the leaflet surface. In the present study, valve canting disrupted this conventional sinus vortex structure, altered the effective available area of both the native sinus and neo-sinus, and redirected the ascending aortic main jet. These changes impaired sinus washout and reduced transport-mediated clearance, particularly within the cusp experiencing the greatest spatial restriction.The resulting low-velocity and recirculatory zones prolonged local residence time near the leaflet surface, creating prothrombotic microenvironments favorable for platelet adhesion and fibrin accumulation. Once platelets adhere to the bioprosthetic leaflet surface, they can provide a procoagulant surface that supports local thrombin generation and fibrin deposition, consistent with the leaflet-associated thrombus identified by Carstairs staining. Under conditions of impaired washout, prolonged platelet residence within recirculating sinus flow likely increases platelet–surface interactions and promotes localized aggregation. These transport-mediated processes provide a plausible explanation for the spatial colocalization between regions of impaired sinus washout and Carstairs-positive platelet and fibrin deposition observed in our experiments. Notably, thrombus formation in our model corresponded more closely with impaired washout than with low instantaneous velocity alone, highlighting the importance of transport processes, particularly recirculation, residence time, and particle trapping, in governing thrombogenic risk. Together, these observations suggest that THV thrombosis may be driven primarily by sinus-resolved transport dynamics rather than by bulk hemodynamic parameters alone.

Clinical imaging studies have previously linked non-coaxial THV deployment and frame deformation with increased incidence of hypoattenuating leaflet thickening (HALT) and leaflet thrombosis.^14,24^ Furthermore, prior benchtop investigations of neo-sinus stasis have shown that even modest deviations from coaxial alignment can lead to changes in flow in the vicinity of THV. However, the mechanistic hemodynamic basis linking such altered flow dynamics and thrombosis has remained uncertain. The present study provides direct experimental evidence linking valve canting to sinus-specific flow disturbances and spatially matched leaflet thrombus deposition. Our results, therefore, while complementing the previous studies, provide a mechanistic explanation for previously reported associations between THV deployment configurations and leaflet thrombosis observed in retrospective clinical imaging studies.

From a clinical and procedural standpoint, these findings emphasize the importance of achieving coaxial alignment during valve replacement procedures. Attention to valve orientation through pre-procedural planning, device positioning, and intraprocedural imaging may reduce unfavorable flow micro-environments and thereby decrease the substrate for leaflet thrombosis and HALT. More broadly, the results highlight the potential value of sinus-resolved hemodynamic assessment, rather than solely observing global valve performance metrics, for identifying patients at risk of post-procedural leaflet thrombosis. Improved understanding of sinus-scale flow transport may also inform future valve design strategies to enhance sinus washout and potentially minimize thrombogenic flow conditions.

This integrated benchtop study has several important limitations. First, experiments were conducted in a single statistically derived “average” aortic root geometry rather than across a range of patient-specific anatomies. Although statistical shape modeling reduces reliance on idealized geometries, anatomic variability may influence in vivo sinus flow patterns and canting effects. Second, the thrombogenicity experiments captured acute thrombus formation in a porcine whole-blood loop and therefore do not fully represent chronic processes leading to organized leaflet thickening or structural valve degeneration. Third, the in vitro system cannot fully replicate in vivo biological factors such as endothelial responses, systemic inflammation, and ventricular-aortic coupling. Finally, each canting condition was evaluated in a single experiment in each modality (PIV and whole-blood loop), yielding 6 total experiments across the study; however, the absence of independent replications within each condition limits formal statistical inference. Nevertheless, the agreement between the PIV-derived flow disturbances and the spatially matched histologic thrombus deposition supports the proposed mechanistic link between canting-induced stasis and leaflet thrombosis.

## 5 Conclusions

In summary, modest deviations from coaxial transcatheter valve deployment can substantially reorganize sinus flow dynamics, creating cusp-specific regions of stagnation that promote leaflet surface thrombosis despite similar global valve hemodynamics across the deployment configurations. These findings highlight the sensitivity of sinus transport dynamics to deployment configurations and highlight the importance of valve alignment in determining post-procedural thrombogenic risk. By integrating high-resolution PIV with a physiologic blood loop and leaflet histology, we directly relate valve canting to sinus flow dynamics, washout, and surface thrombus formation. Future studies may extend these findings to patient-specific root anatomies, investigate surface or pharmacologic strategies to reduce canting-induced thrombosis, and track the evolution of acute thrombus toward imaging-detectable HALT and structural valve degeneration.

## Acknowledgments

This work was supported by the National Institutes of Health, National Heart, Lung, and Blood Institute under award number R01HL167442. The content is solely the responsibility of the authors and does not necessarily represent the official views of the National Institutes of Health. The authors thank Aqua Asberry and the Petit Institute for Bioengineering and Bioscience (IBB) Core Facility, Georgia Tech., for assistance with tissue processing and histology sample preparation.

## Supplemental Figures

**Figure S1:**
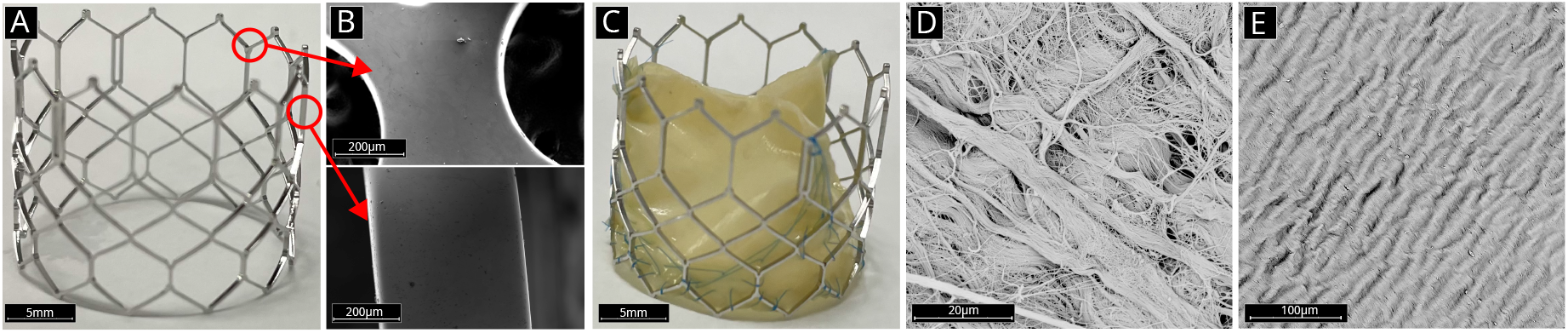
Representative images of a custom transcatheter valve fabrication. (A) Laser-cut Cobalt-Chromium frame prior to leaflet mounting. (B) High-magnification view of representative (red outlines) stent-strut edge demonstrating smooth, burr-free surface after electropolishing. (C) Assembled valve consisting of the Cobalt-Chromium frame and trileaflet porcine pericardial tissue. (D–E) Scanning electron microscopy (SEM) of leaflet microstructure reveals a dense collagen fiber network, i.e., fibrous side facing the ventricle (D) and serous side facing the aorta in (E), consistent with the microstructure of glutaraldehyde-treated pericardial bioprosthetic valve leaflets reported in prior studies.

**Figure S2:**
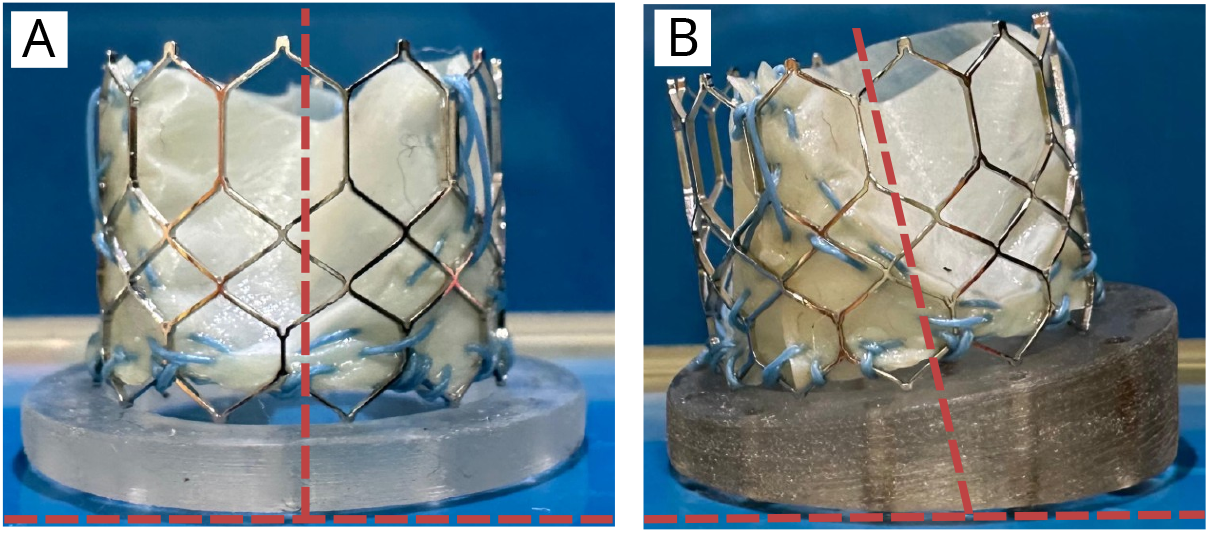
Representative images of in-house THVs mounted over 3D-printed tapered disks to precisely maintain the canting angle.On the left, a control valve with no canting is shown, while on the right, a tapered disk at –10 degrees is shown.

**Figure S3:**
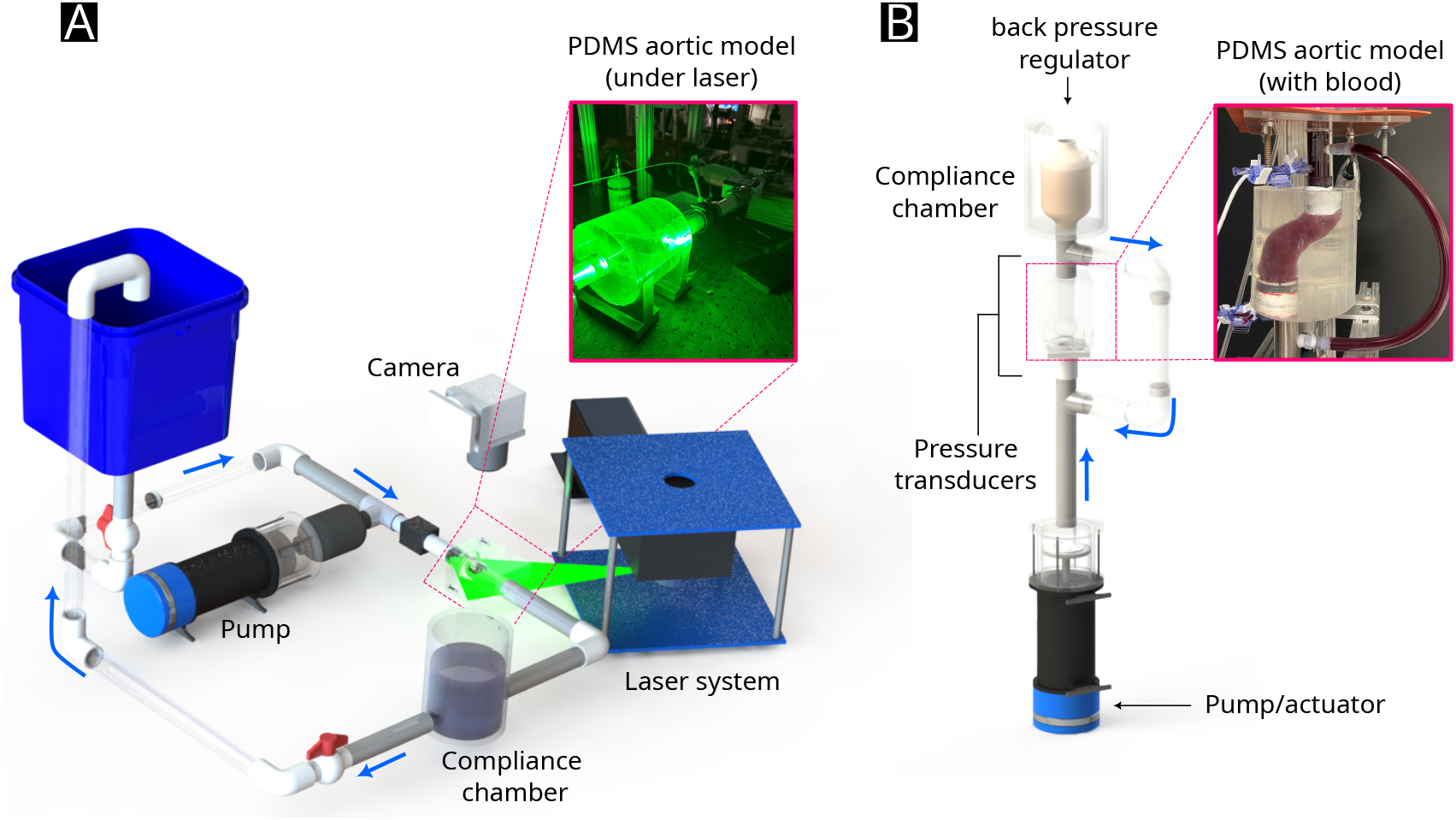
Experimental platforms for hydrodynamic and biological assessment using a population-derived PDMS aortic model. (A) Optical flow loop used for phase-averaged particle image velocimetry (PIV). A pulsatile pump generated physiologic waveforms through a compliance chamber and mechanical mitral valve into the PDMS aortic root model. Flow rate was monitored with an inline probe, and velocity fields were acquired using a laser sheet and high-speed camera (inset). The working fluid was a refractive–index–matched glycerin solution. (B) Blood loop configuration using the same PDMS aortic model. A linear piston actuator generated matched pulsatile flow, with upstream and downstream pressure transducers for hemodynamic monitoring. A compliance chamber and adjustable back-pressure regulator maintained physiologic loading conditions. The transparent PDMS model was perfused with porcine blood for biological assays (inset).

**Figure S4:**
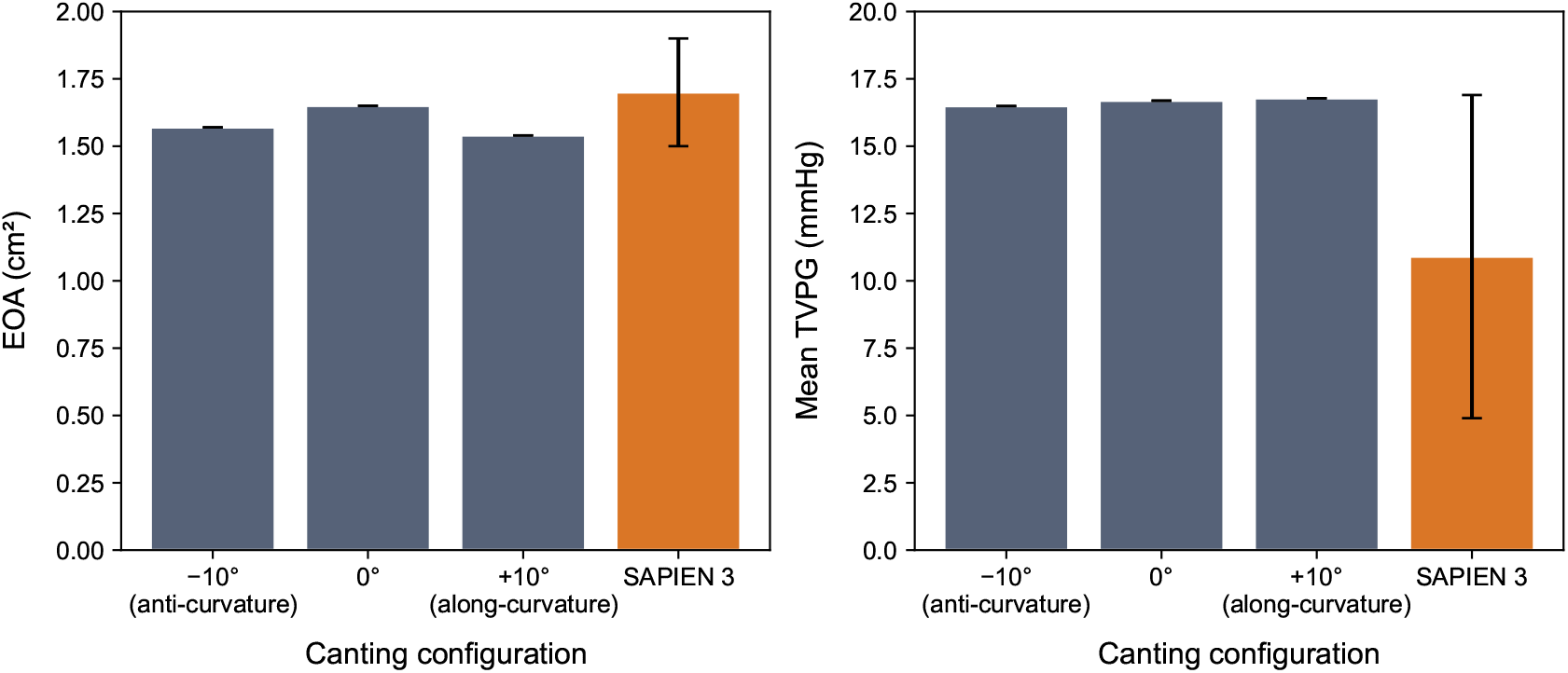
Global hemodynamic performance across canting conditions. Effective orifice area (A) and mean transvalvular pressure gradient (B) remained equivalent across all canting angles and matched published SAPIEN 3 performance [Külling et al. Eur Heart Jou. 2018]

